# Direct visualisation of post-replication gap formation at the bacterial RRS

**DOI:** 10.64898/2026.06.16.732529

**Authors:** Nicholas Kusi-Appauh, Phuong Pham, Elise M. Wilkinson, Michael M. Cox, Jacob S. Lewis, Myron F. Goodman, Lisanne M. Spenkelink

## Abstract

Replication risk sequences (RRS) are recently discovered genomic structural elements that trigger post-replication gap formation during replisome passage. In *E. coli*, the two RRS elements are nearly perfect 222-bp G-quadruplex-containing repeats that flank the terminal domain and are highly conserved in both sequence and genomic position across enterobacteria. We report here the first direct visualisation of RRS function *in vitro* using single-molecule methods. When the G4 strand of the RRS element is positioned on the lagging-strand template, gaps are formed essentially every time a replisome encounters it. An increase in ssDNA in the synthesised DNA is readily seen using ssGAP-seq methods. When the G-quadruplex strand of the RRS is positioned on the leading-strand template, gaps are formed, albeit at lower frequency. However, the continued DNA synthesis in a rolling-circle assay indicates that the gaps are still formed on the lagging strand, indicating that the RRS complementary strand has a significant but reduced capacity to form a structure that triggers lesion skipping. The results document the potency of the RRS as a trigger for gap formation, suggesting a possible function for at least some eukaryotic G-quadruplexes.

## Main

The genomes of almost all organisms contain sequences capable of forming complex intramolecular structures through base-pairing. Many such sequences act at the RNA level, folding within transcripts or encoding structured functional RNAs such as rRNA, tRNA and small RNAs. Others can fold as single-stranded DNA. Prominent examples are G-rich sequences that form G-quadruplexes (G4s), with about 700,000 potential G4-forming sequences identified in the human genome ^1–7^. Many candidate G4s occur in enhancers and promoters ^8^, where regulatory functions are known or likely, but the functions of many others remain unknown. Bacterial genomes typically contain a few dozen to a few hundred G4s ^2^.

We recently developed a new method to examine protein binding to single-stranded DNA on a genomic scale, designated ssGAP-seq ^9^. In applying it to RecA and single-stranded DNA-binding protein (SSB) binding to single-stranded DNA (ssDNA), we identified two sites in the *Escherichia coli* (*E. coli*) genome where both proteins were deposited at elevated rates on the lagging-strand template. At the ends of both sites, we identified a pair of 222-bp GC-rich repeats. These are positioned symmetrically about the terminal *dif* site, each 650-kbp distant from *dif*. In the genome, the RRS elements trigger post-replication gap formation as replisomes pass. We have therefore named them Replication Risk Sequences or RRS ^9,10^. The RRS elements include a G4 on one strand (RRS-G4), which impedes DNA synthesis by *E. coli* DNA polymerases I, II, III, and IV *in vitro* and *in vivo* ^10^.

The generation of programmed single-stranded gaps represents a novel function for a genomic sequence element. The RRS elements are highly conserved in both positioning and sequence among the enterobacteria ^9,10^, although some species have one or more additional RRS within their genomes. Based on their properties *in vivo*, we have postulated that these sequences serve as topological relief valves, with the transient single-stranded gaps helping to accommodate the topological stress accompanying replication termination and chromosome segregation ^10^. Indeed, deletion or replacement of RRS elements in the *E. coli* genome has dramatic effects on global genome topology ^10^. These observations raise the possibility that some eukaryotic G4s may similarly trigger transient gaps during replication at positions that facilitate chromosome function or folding. Only one other bacterial genomic element has been described that promotes gap formation upon replisome encounter: a G4-forming sequence required for pilin antigenic variation in *Neisseria gonorrhoeae* ^11,12^. Identifying genomic sequences with this activity is non-trivial, and additional examples may have been missed.

Although RRS-facilitated gap formation is clear *in vivo*, little is known about how these sequences act mechanistically. Key questions remain: Do replisomes pause when they encounter an RRS? Are other proteins or enzymes required to facilitate bypass of an RRS or is the basic replisome sufficient? How efficient is an RRS in inducing gap formation; i.e., does gap formation occur every time an RRS is encountered by a replisome? Secondary structure calculations show that the complementary strand (RRS-C) can also form intramolecular secondary structure, despite lacking the G4 (Extended Data Fig. 1). Can the RRS-C function as a trigger for gap formation? Will the RRS trigger gap formation if the RRS-G4 strand is incorporated into the leading-strand template? These kinds of questions can only be answered *in vitro*.

To address these and other questions, we have incorporated an RRS into an *in vitro* single-molecule rolling-circle assay ^13,14^. We visualised gaps directly using fluorescently labelled SSB and further analysed the replicated DNA using ssGAP-seq. This approach provides the first direct visualisation of RRS function and reveals a conserved genomic element that efficiently triggers gap formation when encountered by a functional *E. coli* replisome.

### RRS triggers ssDNA gaps *in vitro*

To determine whether the RRS is sufficient to trigger ssDNA gap formation during replication by a purified replisome, we used a single-molecule rolling-circle DNA-replication assay with the minimal E. coli replisome (Fig.1a) ^13,15,16^. Rolling-circle templates containing a single RRS were generated from plasmid DNA using nCas9 ^17^, , allowing us to position the RRS-G4 strand on either the lagging-strand template (RRS-G4-lag; Fig. 1b) or the leading-strand template (RRS-G4-lead; Fig. 1c). We immobilised these templates on a microscope coverslip within a microfluidic flow cell and initiated replication by introducing a laminar flow of buffer containing the components required for leading- and lagging-strand synthesis (Fig. 1a). The growing dsDNA product was stretched by buffer flow and visualised in real time by near-TIRF fluorescence imaging of stained dsDNA, while ssDNA gaps were visualised using fluorescently labelled SSB (Fig. 1a) ^18^.

**Fig. 1:**
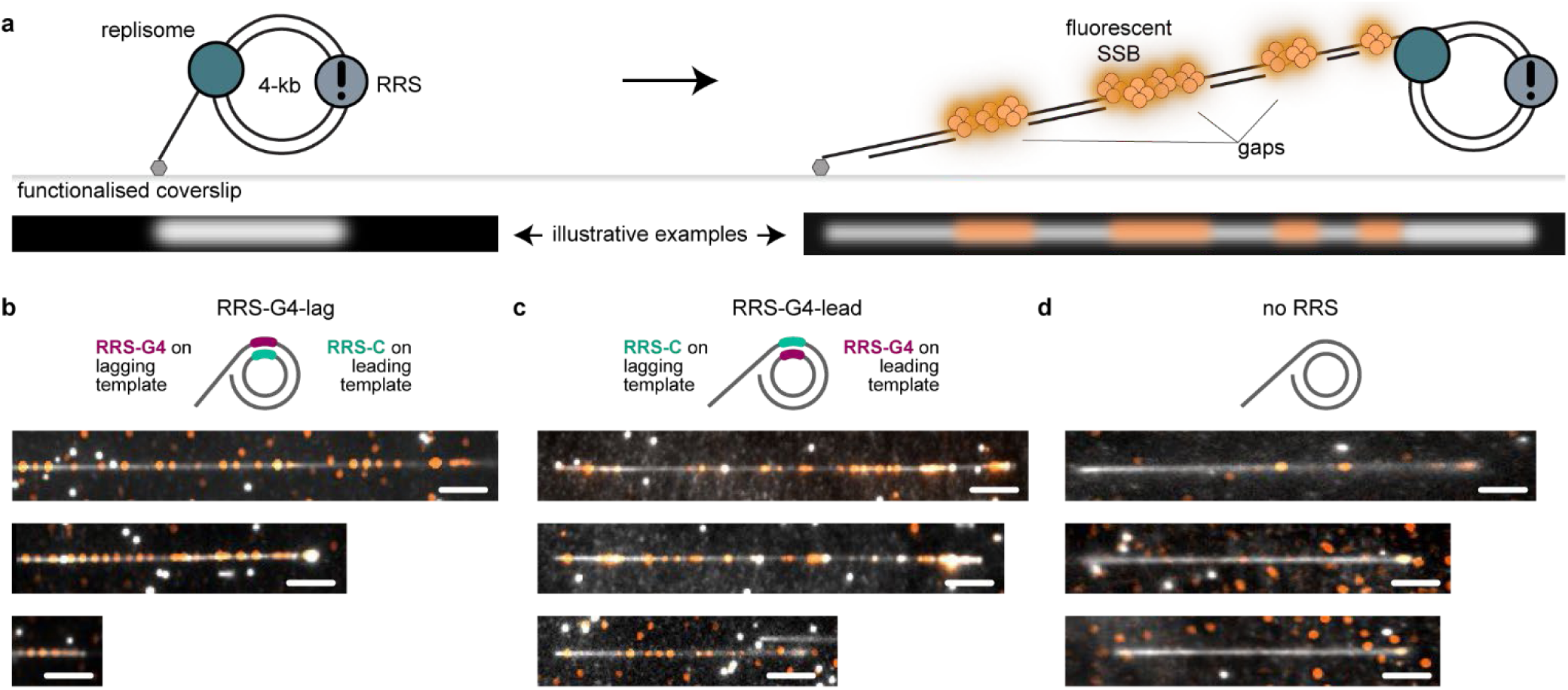
Single-molecule visualisation of rolling-circle replication with an RRS-containing template. **a**, Schematic representation of the rolling-circle DNA replication assay using an RRS-containing DNA template. RRS-containing DNA templates tethered to a functionalised coverslip. Addition of the replication proteins and nucleotides initiates replisome assembly (teal) and DNA synthesis. The DNA products are elongated hydrodynamically by flow, labelled with a fluorescent intercalating DNA stain, and visualised on a TIRF microscope. ssDNA gaps are visualised through fluorescently labelled SSB (orange). **b**, Representative replication products from an assay using the RRS-G4-lag template, show frequent ssDNA gaps (orange) in the dsDNA product (grey). **c**, Use of the RRS-G4-lead also results in frequent gap formation. **d**, Use of a control template without an RRS sequence only shows sporadic gap formation.

We observed frequent ssDNA gaps in assays using the RRS-G4-lag template (Fig. 1b), consistent with *in vivo* observations ^9,10^. Interestingly, frequent gaps also occurred in assays with the RRS-G4-lead (Fig. 1c). In contrast, assays carried out on templates lacking an RRS showed only sporadic gaps (Fig. 1d). Because a leading-strand gap would terminate rolling-circle replication, the continued formation of long replication products suggested that the observed gaps arose predominantly on the lagging strand. We tested this directly using ssGap-seq.

### RRS gaps form in the lagging strand

To determine whether gaps form during leading- or lagging-strand synthesis, we mapped ssDNA in bulk rolling-circle replication products using ssGap-seq ^9,19^. ssGap-seq exploits the ability of sodium bisulfite to convert cytosines to uracils specifically in ssDNA under non-denaturing conditions. Rolling-circle replication was carried out in bulk, after which the replication products were subjected to bisulfite treatment, fragmentation and sequencing to map the location of ssDNA gaps (Fig. 2a).

**Fig. 2:**
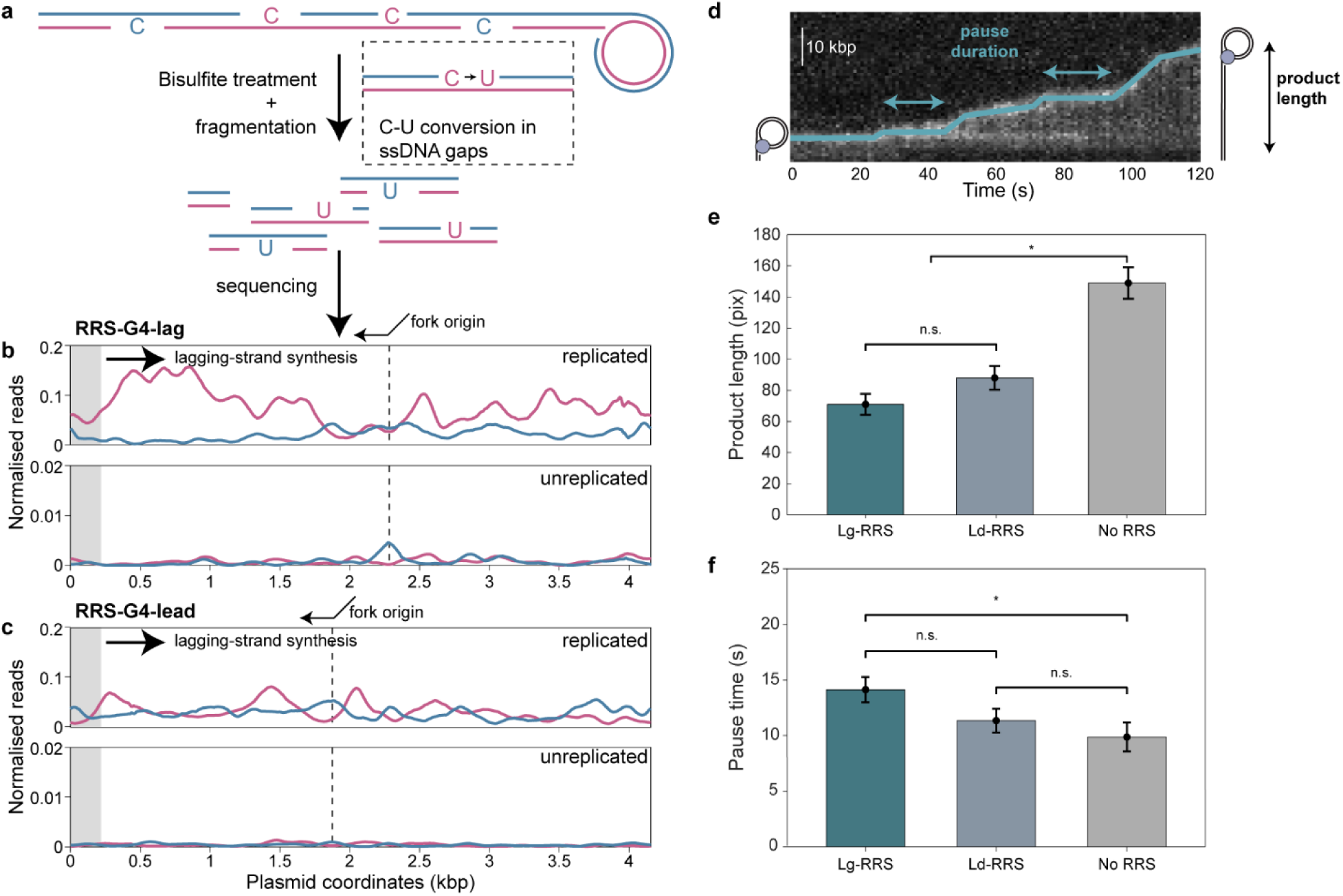
RRS-induced gaps form predominantly in the lagging strand. **a**, Schematic representation of the Gap-Seq workflow. Rolling-circle replication was carried out in a test tube (see methods). Products were treated with sodium bisulfite before fragmentation and sequencing. **b**, Distributions of ssDNA reads on the lagging (purple) and leading (blue) strands of replicated (top) and unreplicated (bottom) RRS-G4-lag. Each data point represents a moving average of ssDNA in a 100 bp window. The position of the RRS sequence is indicated by the shaded grey area. The position and direction of the replication fork are indicated by the vertical line and arrow. The direction of lagging-strand synthesis is indicated by the black arrow. **c**, Gap-seq distributions from reactions using RRS-G4-lead. **d,** Example kymograph showing replication of an RRS-containing template. Tracking of the position of the tip of the DNA, allows identification of individual rate segments (teal). Pause duration is the duration in time where the rate <100 bp/s. The final product length is measured at the end of the experiment. **e,** Mean product lengths from reactions using the RRS-G4-lag (teal), RRS-G4-lead (blue), and a template without an RRS (grey). See also Extended Data Fig. 3. **f,** Mean pause times for replisomes replicating the RRS-G4-lag (teal), RRS-G4-lead (blue) and control template (grey). See also Extended Data Fig. 4). The mean pause durations (± SEM; error bars represent the SEM) are 14.1 ± 1.1 s for RRS-G4-lag (n = 156), 11.4 ± 1.1 s for RRS-G4-lead (n = 113), and 9.87 ± 1.3 s for the control template (n = 57). Statistical comparisons were performed using a likelihood-ratio test for equality of exponential rate parameters, with the significance evaluated using Wilks’ theorem. A p-value <0.05 is denoted by *, and n.s. indicates >0.05. Sample sizes (n) are shown underneath each template.

For replicated RRS-G4-lag templates, most ssDNA reads mapped to the lagging-strand template (∼8%), with a smaller fraction mapping to the leading-strand template (∼2%). Leading-strand ssDNA was distributed relatively uniformly across the template, whereas lagging-strand ssDNA reads were enriched 2–3.5-fold across a 1.2-kb region immediately downstream of the RRS (Fig. 2b, top). This profile indicates frequent lagging-strand gap formation after replisome passage through the RRS. As a control, we measured ssDNA on unreplicated templates. As expected, unreplicated RRS-G4-lag templates showed much lower ssDNA levels on both strands overall (0.07–0.08%), with the highest residual signal localised near the biotin-flap region used to initiate replication (Fig. 2b, bottom).

RRS-G4-lead templates produced a different ssGap-seq profile. ssDNA reads were present at similar levels on the leading and lagging strands across most of the template (∼3% each), with a modest excess of leading-strand reads in the 0.6-kb region downstream of the RRS (Fig. 2c). However, this leading-strand signal does not necessarily indicate frequent leading-strand gap formation. Unlike the single-molecule assay, ssGap-seq measures ssDNA across all molecules in the bulk reaction, including unreplicated and partially replicated substrates. In addition, if the leading-strand polymerase stalls at RRS-G4, continued unwinding by DnaB^20^ could release ssDNA circles that would be detected as leading-strand ssDNA in the ssGAP-seq assay.

Consistent with this interpretation, long replication products were observed for both RRS-G4-lag and RRS-G4-lead templates in the single-molecule assay, whereas leading-strand gap formation would be expected to terminate rolling-circle replication (Extended Data Fig. 2). Product lengths were only modestly reduced for both RRS-containing templates relative to the no-RRS control (Fig. 2d,e; Extended Data Fig. 3), corresponding to leading-strand gap formation in only ∼5% of RRS encounters.

Replication pause analysis further supported the conclusion that leading-strand synthesis is rarely disrupted by the RRS. Replication pausing is an inherent feature of the minimal replisome; on templates lacking an RRS, we observed ∼1.3 pauses per DNA molecule, with a mean pause duration of ∼10 s (Fig. 2f). Pause durations were only modestly increased, to ∼14 s for RRS-G4-lag and ∼11 s for RRS-G4-lead (Fig. 2f; Extended Data Fig. 4).

This small effect on leading-strand synthesis can be rationalised by replisome geometry. The ssDNA between DnaB and the leading-strand polymerase is likely too short or short-lived for secondary structures such as G4s to form efficiently. This interpretation is consistent with previous observations that SSB is not required for efficient leading-strand synthesis, suggesting that the leading-strand template remains largely unstructured ^21,22^, and with single-molecule studies of the T7 replication system ^23^. Thus, the RRS-G4 does not appear to be a major block to leading-strand synthesis.

In this assay, pauses reflect both leading-strand progression and repriming efficiency. Because repriming efficiency is primarily governed by primase ^24–26^, which was held constant across assays, the small differences in pause duration most likely reflect modest differences in leading-strand progression. In the RRS-G4-lag configuration, the leading-strand polymerase encounters RRS-C, whereas the helicase must negotiate RRS-G4 directly, suggesting that RRS-G4 is a mild obstacle to helicase progression.

### RRS orientation controls gap frequency and size

Having established that the RRS predominantly affects lagging-strand synthesis, we next determined how efficiently it induces gaps. First, we measured the distance between successive gaps (Fig. 3a,b). For the RRS-G4-lag, the characteristic inter-gap distance was ∼4 kbp (Fig. 3c, Extended Data Fig. 5). This distance matches the size of the rolling-circle template, indicating that a gap is created every time the replisome encounters the RRS. For the RRS-G4-lead, this distance was ∼6 kbp, corresponding to a gap-creation frequency of ∼66% (Fig. 3c, Extended Data Fig. 5).

**Fig. 3:**
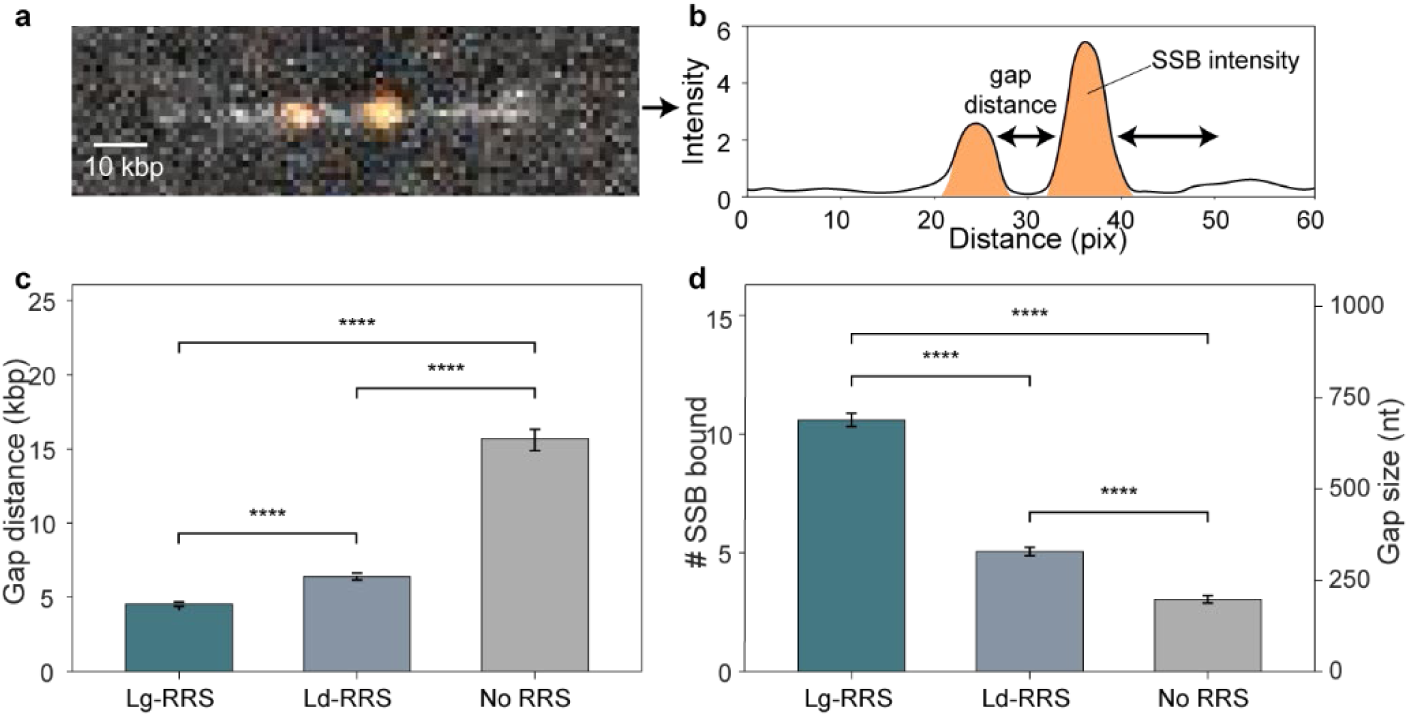
Quantification of RRS-induced gap frequency and size. **a**, Representative replication product from a reaction on an RRS-containing DNA template. **b**, The fluorescence intensity of the SSB-coated ssDNA gaps in (a) is used to calculate the distance between gaps and the number of SSBs bound. (**c**) Distances between the ssDNA gaps in the replicated RRS-G4-lag (teal), RRS-G4-lead (blue), and control template (grey). See also Extended Data Fig. 5). The characteristic distances (± SEM; error bars represent the SEM) are 4.54 ± 0.14 kbp for RRS-G4-lag (n = 1,036), 6.40 ± 0.24 kbp for RRS-G4-lead (n = 716), and 15.62 ± 0.78 kbp for the control template (n = 403). **d**, Size of ssDNA gaps in replicated RRS-G4-lag (teal), RRS-G4-lead (blue), and control templates (grey). See also Extended Data Fig. 6. The mean gap sizes (± SEM, error bars represent the SEM) are 698.4 ± 18.2 nt for RRS-G4-lag (n = 1,415), 328.3 ± 11.7 nt for RRS-G4-lead (n = 810), and 197.6 ± 10.4 nt (n = 386) for the control template. Statistical comparisons of all exponential distributions are carried out using a likelihood-ratio test for equality of exponential rate parameters, with the significance evaluated using Wilks’ theorem (p < 0.0001 for all pairwise comparisons, ****). Sample sizes (n) are shown underneath each template.

We also measured gap size by quantifying the integrated SSB intensity at each gap (Fig. 3b, orange). Dividing this intensity by the intensity of a single SSB molecule gives the number of SSBs bound per gap (Fig. 3d, Extended Data Fig. 6). Approximately 10 SSBs bound per gap in RRS-G4-lag assays and ∼5 in RRS-G4-lead assays. Assuming SSB binds in the 65-nt mode ^27,28^, corresponds to gap sizes of ∼650 nt and ∼325 nt, respectively. These values agree with the sizes we obtained through duplex DNAse treatment (Extended Data Fig. 7) and are similar to those observed *in vivo* ^9,10^.

Gap size is determined by the frequency of priming and the ability of the lagging-strand polymerase to synthesise through the RRS. As priming efficiency is primarily governed by DnaG and SSB concentrations ^15,29^), which are the same in all our assays, differences in gap size between templates must reflect differences in the ability of the lagging-strand polymerase to synthesise through the RRS. The higher gap frequency and larger gap size in RRS-G4-lag assays indicate that the RRS-G4-lag template more effectively disrupts lagging-strand synthesis than the RRS-G4-lead template. This observation shows that the RRS-C is a weaker block to the polymerase.

## Discussion

The work presented here has three principal conclusions. First, RRS elements are sufficient to program post-replication gap formation during replisome passage, with the native RRS orientation, in which RRS-G4 is copied as the lagging-strand template, triggering gap formation at nearly every replisome encounter. Second, the complementary strand, RRS-C, can also trigger gap formation, albeit with reduced efficiency. Third, neither RRS strand efficiently promotes gap formation when copied as the leading-strand template, consistent with a model in which RRS activity depends on transient exposure of the lagging-strand template during Okazaki-fragment synthesis.

Based on these observations, we propose a strand-exposure model for RRS-triggered gap formation (Fig. 4). During each Okazaki-fragment cycle, the lagging-strand template is transiently single-stranded, allowing the RRS to fold. When RRS-G4 is on the lagging-strand template, lagging-strand synthesis is blocked by the G4 and a gap is generated in nearly every replisome encounter (Fig. 4, top). When RRS-C is on the lagging-strand template, the structure is a weaker block, and gap formation occurs in approximately two-thirds of encounters (Fig. 4, bottom). By contrast, the close proximity of DnaB and the leading-strand polymerase likely limits the lifetime of ssDNA on the leading-strand template, reducing the opportunity for RRS folding and allowing leading-strand synthesis to continue (Fig. 4).

**Fig. 4.**
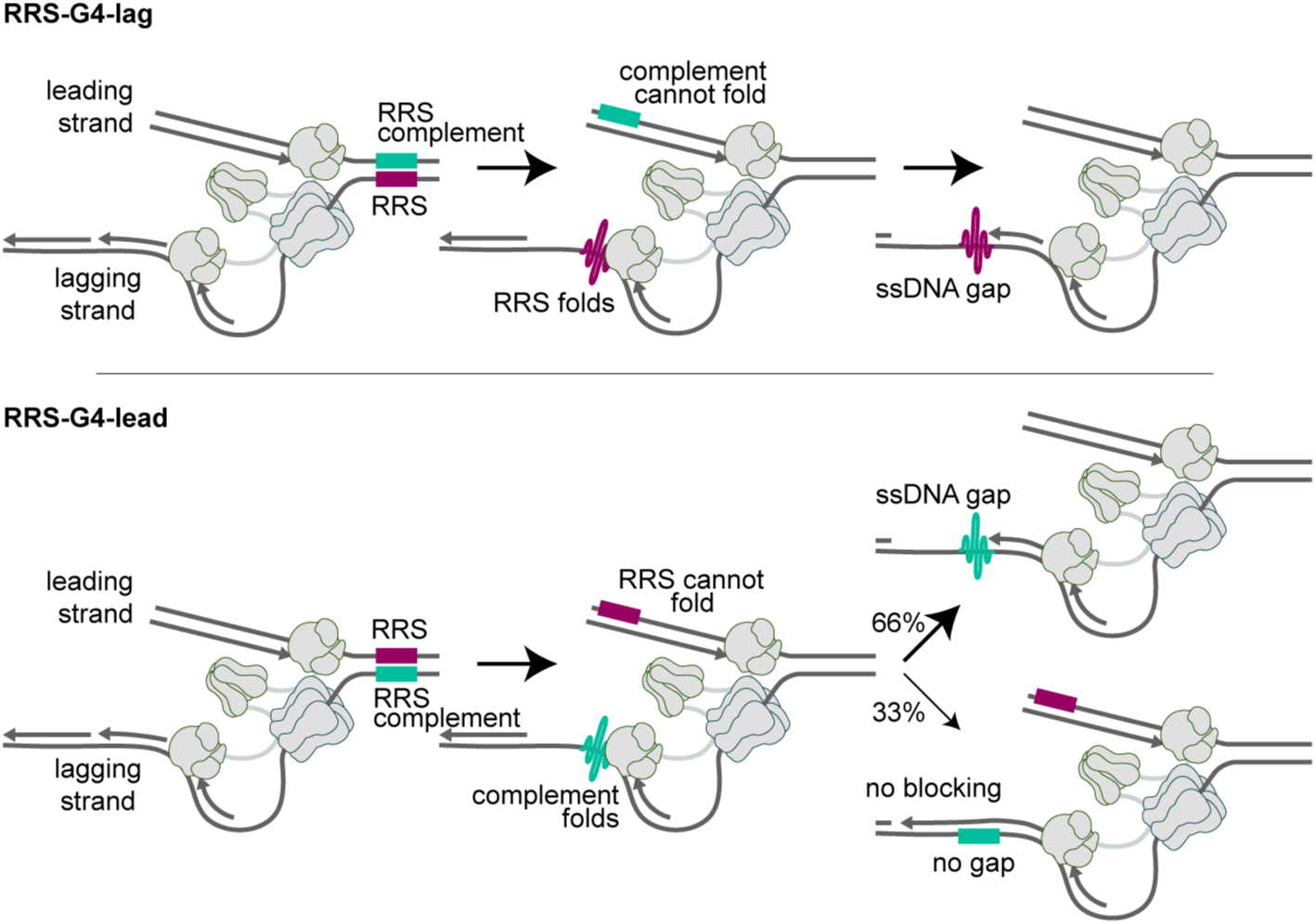
Mechanism for post-replicative gap formation at an RRS. (top) Schematic showing replication of DNA with the RRS-G4 (purple) on the lagging-strand template. Following unwinding of the parental DNA by the helicase, the lagging-strand template is transiently single stranded because of the discontinuous nature of Okazaki-fragment synthesis. This allows the RRS-G4 to fold into its secondary structure. Lagging-strand synthesis is blocked by the RRS-G4, resulting in an ssDNA gap corresponding to the size of the unfinished Okazaki fragment. (bottom) Schematic showing replication of DNA with the RRS-G4 on the leading-strand template. In this orientation, the RRS-C (green) secondary structure forms on the lagging-strand template. This structure is a weaker block to the polymerase, blocking lagging-strand synthesis in only ∼66% of encounters. (top and bottom) Due to the spatial arrangement of the leading-strand polymerase relative to the helicase, neither the RRS-G4 nor the RRS-C can fold into their secondary structures when present on the leading-strand template.

A programmed gap in genomic DNA is a problem after chromosome segregation. If the gap is not closed, the next replication cycle would create a double strand break. Therefore, RRS-induced gaps must be transient. Gap filling is likely to be challenging because the folded RRS-G4 is a strong block to DNA synthesis: cloned RRS elements cannot be sequenced by standard commercial sequencing unless additional heat is used ^10^. Filling these gaps may therefore require specialised polymerase and helicase activities. RecQ and/or Rep helicases are plausible candidates ^30,31^.

More broadly, our findings establish programmed ssDNA-gap formation as a potential function of evolved and conserved genomic sequence elements. Only two examples of such elements have now been described, both in bacteria. This may suggest the phenomenon is rare. However, the phenomenon may be much more prevalent in genomes of many organisms. Recognition that such elements exist has had two barriers. First, their existence was unexpected. Second, the methodology needed to detect them has only recently become available. Elucidation of the function of the G4- forming structure that triggers gap formation in pilin antigenic variation in *Neisseria gonorrhoeae* required many years of careful investigation ^11,12^. For a decade or so, it remained the only known example. Discovery of the RRS elements was a serendipitous outcome of the development of ssGAP-seq ^9^. Transient ssDNA can relieve topological stress by providing rotational flexibility, with every phosphodiester bond acting as a functional swivel. Foldable DNA sequences may therefore serve as replication-coupled topological release valves. In principle, any DNA sequence that can fold up quickly — a G4 or any other stable tertiary structure — could potentially block full replication of the lagging-strand template and create a transient gap. A key future question is whether some eukaryotic G4s or other foldable sequences similarly promote transient gap formation at sites that contribute to chromosome organisation, chromosome folding, or replication-associated structural transitions.

## Materials and Methods

### Replication proteins

The *E. coli* DNA replication proteins were produced as described previously: the DnaB_6_(DnaC)_6_ helicase–loader complex ^32^, β_2_ sliding clamp ^33^, DnaG primase ^34^, the Pol III τ_3_δδ′ψχ clamp loader ^35^ and Pol III αεθ core ^14,36^. The SSB-K43C was expressed, purified and labelled with AlexaFluor-647 as previously described ^18^.

### Construction of rolling-circle replication templates

#### Custom DNA & RNA oligonucleotides

The DNA oligonucleotides, CRISPR RNA’s (crRNA) and Trans-activating CRISPR RNA (tracrRNA) used in template construction were purchased from Integrated DNA Technologies (IDT). The sequences of these oligos are listed in Extended Data Table 1. All DNA and RNA oligonucleotides were stored in TE buffer (10 mM Tris-HCl pH 8.0, 1 mM EDTA) at –80 °C.

#### Transforming, culturing and purifying RRS-containing plasmids

The *lysO* RRS, spanning the bases 168-389 upstream of the *lysO* gene, were cloned into the 4213 bp plasmid EAW1307 (pEAW1307). This was achieved by cloning the *lysO*-RRS sequence into the EcoRI and BamHI sites of the 4361 bp pBR322, a common plasmid cloning vector.

A smaller, 2252 bp RRS-containing plasmid was cloned into the smaller 2030 bp pSCW01 (Addgene plasmid #72300) by the insertion and ligation of the *lysO*-RRS gene sequence (Aldevron, USA). The 2.2 kbp plasmid (pNKA001) was first resuspended to 50 ng/µL in TE buffer. Then, 75 ng plasmid was added to 50 µL 10β *E. coli* cells and left to incubate on ice for 30 min. The solution was then heat shocked at 42 °C for 30 s and then left on ice for 5 min. Following this, 200 µL SOC outgrowth medium was added, and the solution was incubated at 37 °C for 1 h with 900 rpm rotation. This solution was plated onto LB agar plates supplemented with 100 µg/mL ampicillin and left to incubate overnight at 37 °C.

Both the 4.2 kbp pEAW1307 and 2.2 kbp pNKA001 needed to be cultured in bacteria to permit the rapid duplication of the plasmid for template construction. Bacterial cultures were initiated by inoculation from the pEAW1307 glycerol stock or by isolating individual colonies from the pNKA001 transformation plates. Cells transformed with pEAW1307 or pNKA001 were grown in LB medium (100 µg/mL ampicillin) at 37 °C for 18 h with 180 rpm rotation. The cell cultures were then centrifuged at 4,600 x *g* for 10 min at RT, and the plasmid DNA was extracted using the QIAprep Spin Miniprep Kit, in accordance with the manufacturer’s protocol. The DNA was then resuspended in TE buffer, and the concentration of each purified plasmid was determined by measuring the absorbance at 260 nm using a NanoDrop spectrophotometer.

#### Turning RRS-plasmids into rolling-circle templates

Once the pEAW1307 and pNKA001 DNA was extracted and purified, the next step was to turn both plasmids into a rolling-circle template using two distinct template construction protocols. These dsDNA rolling-circle templates contain a 5′-ssDNA overhang, which serve as the replication fork onto which the replisome can load. When replication begins, the internal (inner) and external (outer) strands act as the templates for leading-strand synthesis and lagging-strand synthesis, respectively. When the template is tethered to the surface of a streptavidin-functionalised coverslip within the confines of a microfluidic flow cell device, this template can be replicated by the activity of the *E. coli* replisome. This will result in the elongation of the replicated dsDNA template in the direction of the laminar buffer flow at a speed determined by the replication rate of the replisome.

To convert both the 2.2 kbp pNKA001 template (“Ld-RRS2”) and the 2 kbp (“control”) pSCW01 into rolling-circle templates, a previously-described protocol was followed ^37^. Briefly, both plasmids were nicked by Nt.*Bst*NBI in four distinct sites on the same strand. Once these nicked oligonucleotides are displaced, a 37 nt-long ssDNA region is generated. A partially complementary fork oligonucleotide can then be annealed and ligated to this gap to create a forked overhang capable of initiating and sustaining replication. The RRS in the Ld-RRS2 template is oriented relative to the fork to ensure the sequence capable of forming the GQ undergoes lagging-strand replication.

The 4.2 kbp pEAW1307 template was converted into a rolling-circle template using an nCas9-based method as previously described ^17^. Briefly, four gRNA complexes are generated by individually hybridising tracrRNA with different crRNA. Then, the pEAW1307 is nicked in four sites in three sequential steps by nCas9 functionalised with the four distinct gRNA complexes. Once nicked, the oligonucleotides are displaced by the addition of an excess of capture oligos, with a partially complementary fork oligonucleotide being annealed and ligated to this gap. This generated replication fork is fundamentally no different from those present on the Ld-RRS2 and control rolling-circle templates. By altering the identity of the four crRNA, the four different nCas9-gRNA complexes can either nick the outer or inner strand of the plasmid. With the addition of the complementary capture oligos and the corresponding fork, the GQ-forming region of the RRS can be tuned to undergo either leading or lagging-strand synthesis, depending on its relative position with the generated replication fork. These rolling-circle templates were dubbed “RRS-G4-lag” and “RRS-G4-lead”. The crRNA, capture oligos and forked primer used to generate RRS-G4-lag and RRS-G4-lead are shown in Extended Data Table 1.

Once the rolling-circle templates were generated, the excess oligonucleotides were removed via size selection with AMPure XP beads (Beckman Coulter Life Sciences). By adding a 0.6:1 ratio of beads:DNA template, the larger rolling-circle templates could be annealed to the beads, allowing for their purification. The purified rolling-circle templates were eluted in TE buffer and stored at -80 °C.

### In vitro single-molecule rolling-circle DNA replication assay

#### In-vitro single-molecule microscopy

The single-molecule microscopy experiments described here were carried out on an Eclipse Ti-E inverted microscope (Nikon, Japan) which is fitted with a CFI Apo TIRF 100X oil immersion objective (NA 1.49, Nikon). The movies were recorded using a Hamamatsu C9100-13 512 x 512-pixel EM-CCD camera (Andor iXon 897). An electrically heated chamber coupled to an objective heating jacket (Okolab, USA) was used to maintain a 33 °C temperature throughout the duration of the experiments. DNA fluorescently stained with SYTOX Orange were imaged with a 514 nm laser (Coherent, Sapphire 514-150 CW) at 420 mW/cm^2^. The AF647-labelled SSB-K43C was visualised and imaged with a 647 nm laser (Coherent, Obis 647-100 CW) at 220 mW/cm^2^. All movies were acquired with an exposure time of 200 ms.

#### Flow cell preparation for in vitro imaging

Microfluidic flow cells needed to be constructed to carry out single-molecule replication experiments. These flow cells consist of a PEG-biotin-functionalised glass cover slips layered by a PDMS chamber as described ^38^. Once the inlet and outlet tubing have been inserted into the PDMS chamber, the flow cell was blocked with Tween-20 blocking buffer (50 mM Tris-HCl pH 7.6, 50 mM KCl, 2% (v/v) Tween- 20) for 15 min to prevent non-specific interactions. Following this, a wash buffer (25 mM Tris-HCl pH 7.6, 10 mM magnesium acetate, 250 mM potassium glutamate, 40 µg/mL BSA, 0.1 mM EDTA, 5 mM DTT, 0.0025% (v/v) Tween-20) to remove the blocking buffer and prepare the flow cell for replication buffers.

#### Single-molecule E. coli DNA rolling-circle polymerases-in-solution replication assay

The rolling-circle template was pre-incubated with 30 nM DnaB_6_(DnaC)_6_ in replication buffer supplemented with 10 mM DTT, 1 mM ATP and 150 nM SxO at 37 °C for 1 min. The DNA-DnaBC solution was subsequently flowed into the flow cell at a rate of 10 µL/min for between 1-10 min, using a DNA concentration of 5-50 pM to achieve ∼50 molecules per FOV. Replication was then initiated by flowing in a solution containing 10 mM DTT, 1.25 mM ATP, 250 µM CTP, GTP and UTP, 50 µM dATP, dCTP, dGTP and dTTP, 30 nM αεθ core, 10 nM τ_3_δδ′ψχ clamp loader, 150 nM DnaG primase, 30 nM β_2_ clamp, 20 nM AF647-labelled SSB-K43C and 150 nM SxO. This was flowed in at 20 µL/min to facilitate the elongation of the rolling-circle templates in the direction of the laminar buffer flow once replication had propagated. During this time, seven fields of view were imaged for 2 min before moving to a new, non-overlapping field of view. Once complete, five fields of view were imaged for 1 min to examine the SSB binding profiles.

#### Single-molecule E. coli DNA rolling-circle pre-assembly replication assay

The rolling-circle template was pre-incubated with 30 nM DnaB_6_(DnaC)_6_ in replication buffer supplemented with 10 mM DTT, 1 mM ATP and 150 nM SxO at 37 °C for 1 min. The DNA-DnaBC solution was subsequently flowed into the flow cell at a rate of 10 µL/min for between 1-10 min, using a DNA concentration of 5-50 pM to achieve ∼50 molecules per FOV. Following on, 30 nM αεθ core and 10 nM τ_3_δδ′ψχ clamp loader were incubated in replication buffer supplemented with 10 mM DTT, 1 mM ATP, 50 µM dCTP, 50 µM dGTP, and 150 nM SxO at 37 °C for 3 min. Once complete, this solution was loaded into the flow cell at 10 µL/min for 10 min. To remove any unbound proteins, a replication buffer “wash” containing 10 mM DTT, 1 mM ATP, 50 µM dCTP, 50 µM dGTP, and 150 nM SxO was flowed into the flow cell at a rate of 200 µL/min for 30 s. Replication was then initiated by flowing in a solution containing 10 mM DTT, 1.25 mM ATP, 250 µM CTP, GTP and UTP, 50 µM dATP, dCTP, dGTP and dTTP, 150 nM DnaG primase, 30 nM β_2_ clamp, 20 nM AF647-labelled SSB-K43C and 150 nM SxO. This was flowed in at 20 µL/min to facilitate the elongation of the rolling-circle templates in the direction of the laminar buffer flow once replication had propagated. During this time, seven FOVs were imaged for 2 min before moving to a new, non-overlapping FOV. Once complete, five FOVs were imaged for 1 min to examine the SSB binding profiles.

### Analysis of the size and frequency of the ssDNA gaps within the replicated templates

All analyses were carried out using ImageJ/Fiji (1.51w) and in-house-built plugins. The intensity of a single SSB monomer was first measured per experiment by first generating an average projection of the 647 channel of the first movie per experiment. By integrating the intensities of all spots present in this channel and fitting a Gaussian distribution to the dataset, the fluorescent intensity of a single SSB monomer could be deduced per experiment. When analysing the 1 min movies at the end of each experiment, which showed SSB binding patterns along the length of the replicated DNA templates, the SSB peaks that colocalised with a replicated DNA template were identified, and their intensities were integrated. The intensity of each SSB spot could then be divided by the calculated intensity of a single SSB monomer to determine the number of SSBs present at each foci. To calculate the distance between each of these SSB foci, which essentially represents the dsDNA bridging the ssDNA gaps, the SSB foci intensities were reproduced as a line plot. Foci over a certain threshold were identified as a gap and were analysed in MATLAB 2021b. Here, the distance between each of the gaps was calculated in pixels, and this could be converted into the length of dsDNA bridging the ssDNA gaps in kilobases.

### Analysis of the replication rate and pause times

To analyse parameters such as replication rate and stalling behaviour, individual DNA molecules were manually selected from each FOV in ImageJ/Fiji. Kymographs, or two-dimensional representations of the individual replication products, which display all frames of the movie arranged along the time axis, were then generated for each selected template. These kymographs facilitate the replication dynamics to be visualised over the course of the acquisition. Individual rate segments were then determined by eye, with a segmented line drawn over the molecule’s tip to track its movement. Using in-house built plugins, the parameters of the drawn lines can be extracted, which can then be used to identify the product lengths, replication rates and stalling events for individual DNA molecules.

### Ensemble DNA rolling-circle replication assay using streptavidin beads

Magnetic streptavidin beads were used to replicate the rolling-circle templates before subsequently isolating the replicated products, which is not achievable within the confines of a microfluidic flow cell device. First, 25 µg T1 MyOne Streptavidin-coupled Dynabeads (Thermofisher, Australia) were washed and equilibrated in 100 µL replication buffer supplemented with 10 mM DTT thrice and then resuspended to 10 µL. Then, 30 nM DnaB_6_(DnaC)_6_ was pre-incubated with 50 ng rolling-circle DNA template in replication buffer supplemented with 10 mM DTT and 1 mM ATP at 37 °C for 5 min. This 40 µL solution was then added to the resuspended beads and allowed to immobilise to the beads with gentle rotation on a HulaMixer at RT for 30 min. By sequestering the beads with a magnet, they were washed three times in 200 µL replication buffer to remove any unbound DNA. Following on, 30 nM αεθ core and 10 nM τ_3_δδ′ψχ clamp loader were incubated in 40 µL replication buffer supplemented with 10 mM DTT, 1 mM ATP, 50 µM dCTP and dGTP at 37 °C for 5 min. This solution was then loaded onto the beads and allowed to incubate at 37 °C for 15 min with mixing at 300 rpm in a Thermocycler to ensure a homogenous bead solution. Once complete, the beads were washed five times in replication buffer to ensure no residual polymerases were present, as these polymerases would fill in potential ssDNA gaps. Replication was initiated by resuspending the beads in 40 µL replication buffer containing 10 mM DTT, 1.25 mM ATP, 250 µM CTP, GTP and UTP, 200 µM dATP, dCTP, dGTP and dTTP, 150 nM DnaG primase, 30 nM β_2_ clamp and 20 nM SSB. Replication was allowed to proceed by incubating the solution at 37 °C for 20 min with 400 rpm rotation. To quench replication, 110 mM NaCl and 50 mM EDTA were added, and the bead solution was boiled at 70 °C for 10 min before returning to RT slowly. The beads were then separated from the solution using a magnet, and the supernatant containing the replicated DNA templates was recovered.

### Mapping ssDNA gaps by *ssGap-seq*

#### Bisulfite treatment of replicated and unreplicated pEAW1307

ssGap-seq to map ssDNA in unreplicated and replicated pEAW1307 was carried as described previously ^9,10^.Non-denaturing bisulfite treatment was carried out for unreplicated or replicated pEAW1307 (10 ng to 50 ng) in the presence of spiked-in 450 ng genomic DNA purified from an E. coli strain EAW1710 lacking both RRS sequences ^10^. To convert C to U deamination on ssDNA gaps, DNA was incubated in freshly prepared solution of 5 M sodium bisulfite and 20 mM hydroquinone (500 ml volume) at 37 °C ^9,10^. Following incubation for 18 h, bisulfite-treated DNAs were washed 3 times with water using Amicon Ultra 10 K Centrifugal Filter Device (Millipore) and desulphonated in 0.3 M NaOH solution for 20 min at room temperature. After removal of NaOH by the Amicon Centrifugal Filter Device, the DNA solution was buffer-exchanged to 10 mM Tris pH 8.0 using Bio-spin P6 column (Bio-Rad, CA).

#### Illumina library preparation and genome sequencing

Purified bisulfite-treated DNA was sheared to an average size of ∼200-250 bp by sonication using a Covaris S2 instrument. Shared DNA was purified and concentrated using a Monarch PCR & DNA “clean-up” kit (New England Biolabs) and quantified by Qubit. 1 ng to 2 ng of sheared DNA was used directly in Illumina’s sequencing library preparation. xGen Methyl-Seq Library kit (Integrated DNA Technologies, IA). The library was prepared according to the company’s protocol with 16 PCR amplification cycles. This library preparation kit relies on enzymatic attachment of adaptors directly on bisulfite-treated ssDNA by the company’s Adaptase technology. Each library was purified using two rounds of an SPRI bead clean up, quantified by Qubit and qPCR, and subjected to Illumina’s sequencing (2 x 150 bp paired-ends) on a Mini-Seq using Mid Output Reagent Kits (300-cycles).

#### Sequencing data analysis

Illumina sequencing data were analysed by CLC Genomic Workbench (v. 21.0.5) (Qiagen). High quality sequencing reads were imported and trimmed 25 bases from the 5’-end and 20 nt from the 3’- end. Sequencing reads having a length of 101 to 105 nt long were used to analyse ssDNA gaps. Bisulfite-treated reads were aligned and mapped to the pEAW1307 reference plasmid using a Bisulfite Sequencing (BS-seq) directional protocol with a length fraction setting of 0.8 and similarity fraction setting of 0.9. The pEAW1307 plasmid sequence was also converted into two different *in silico* versions. For the CT-converted plasmid all C residues were replaced by T, and for the GA-converted plasmid, all G residues were converted to A. To map ssDNA reads to W- and C-strands, reads mapped to pEAW1307 were independently mapped to the CT- and GA-converted plasmids with a length fraction setting of 0.8 and similarity fraction setting of 0.9. The mapped reads were used to analyse the spatial distribution of ssDNA gaps on W- or C- plasmid strands, respectively.

#### Statistical analysis

Mapping coverages with detailed per-base coverage information for BS-seq maps and ssDNA maps for W- and C-strands were exported from the CLC Workbench program in a tab delimited format. Percentages of ssDNA, normalised coverage of ssDNA on W- and C-strands and enrichment were computed using custom-written R scripts.

### Measurement of ssDNA gap sizes in replicated pEAW1307 by duplex DNAse

Measurement of average size of ssDNA gaps was carried out using an engineered dsDNA-specific DNA endonuclease that preferentially degrades dsDNA over ssDNA. Preferent digestion of dsDNA by the duplex DNAse was verified by examining its activity on dsDNA and ssDNA substrates. Duplex-DNAse treatment was carried out by incubating 100 ng of replicated pEAW1307 with 0.2 units of duplex DNAse (New England Biolabs Cat#7635S) in reaction buffer r2.1 (total volume 20 μl). Digestion of dsDNA was carried out at 37°C for 20 min. Undigested ssDNA fragments were purified by Monarch PCR & DNA “clean-up” kit (New England Biolabs) and quantified by Qubit. Purified ssDNA fragments (1 ng to 2 ng) were used as input in xGen Methyl-Seq Library kit (Integrated DNA Technologies, IA) to attach the Illumina’ sequencing adapters at both ends with following modification. i, The enzymatic attachment of the first adaptor directly on ssDNA fragments by Adaptase technology was carried out without preceding denaturation step and with much longer extension time of 15 min at 65°C (standard protocol for extension is 5 min) to allow DNA synthesis completion of long ssDNA fragments; ii, After ligation of the second adapter, AMPure XP-bead purified ligated ssDNA fragments were amplified by PCR for 12 cycles (98°C – 30 s; 60°C - 30 s, 68°C – 10 min) using ^32^P-labelled primers P5 (5’-AATGATACGGCGACCACCGAGATCT-3’ ) and P7 (5’-CAAGCAGAAGACGGCATACGAGAT-3’ ). 5’- end labelling of the primers and 1 kb DNA ladder (New England Biolabs) was carried out using T4 Polynucleotide kinase (New England Biolabs) and ATP[γ-32P] (Revvity). Excess of free ^32^P-ATP in labelled ladder and amplified PCR products was removed using Bio-spin P6 column (Bio-Rad, CA) according to the manufacturer instruction. The cleaned-up PCR products were resolved on a standard 1% agarose gel, visualised on a Typhoon FLA 9500 phosphor-imager, and analysed using Image Quant software. The average ssDNA gap size was determined after subtraction of 130 bp contributed by the Illumina adapters.

## Data availability

Single-molecule and Gap-seq data have been deposited at 10.5281/zenodo.19655914 and are publicly available as of the date of publication.

## Code availability

All analyses were carried out using ImageJ/FIJI (1.51w) ^39,40^ and MATLAB 2016b, and in-house built plugins (https://github.com/Single-molecule-Biophysics-UOW).

## Acknowledgements

We thank members from the Spenkelink & Lewis labs (University of Wollongong), the Keck lab (University of Wisonsin-Madison), the Goodman lab (University of Southern California) and Prof. Antoine van Oijen (University of Sydney), for helpful discussions. Purified *E. coli* DNA replication proteins were a kind gift from Dr Slobodan Jergic, Dr Zhi-Qiang Xu, and Prof. Nicholas E. Dixon (University of Wollongong). pEAW1307 was a kind gift from Dr Elizabeth A. Wood.

## Author contributions

Conceptualisation: N.K-A., P.P., M.F.G., M.M.C., L.M.S.; Methodology, N.K-A., P.P., L.M.S.; Formal analysis: N.K-A., P.P.; Investigation: N.K-A., P.P., E.M.W.; Writing - Original Draft: N.K-A., P.P., M.M.C., L.M.S.; Writing - Review & Editing: E.M.W., M.F.G., J.S.L.; Visualisation: N.K-A., P.P., E.M.W., L.M.S.; Supervision: M.F.G., M.M.C., J.S.L., L.M.S.; Funding acquisition: M.F.G., M.M.C., J.S.L., L.M.S.

## Extended Data

**Extended Data Fig. 1.**
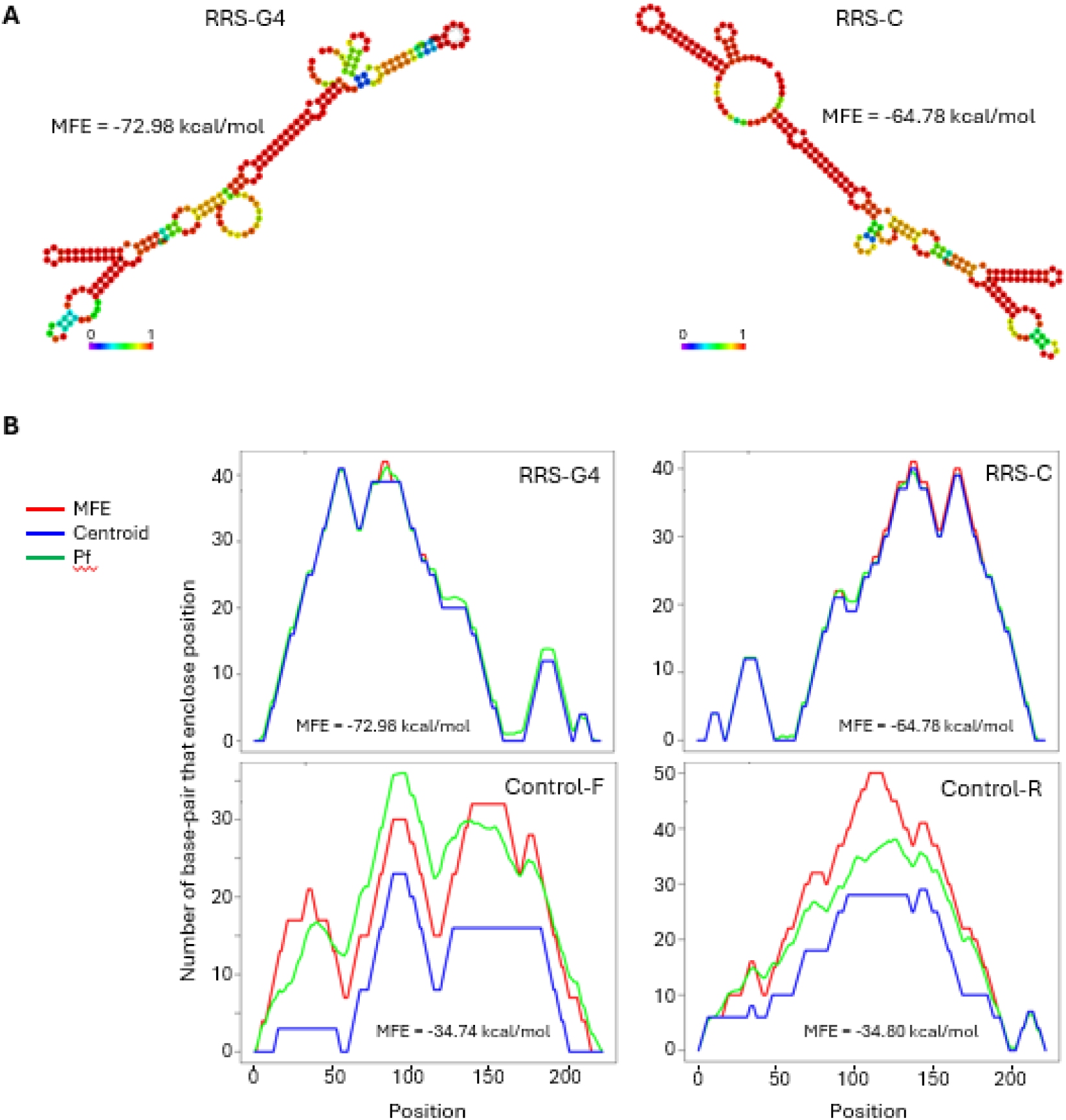
Both RRS-G4 and RRS-C strands can form stable secondary structure when present as ssDNA. **A**, Predicted Minimum Free Energy (MFE) secondary structure with base-pair probabilities for RRS-G4 and RRS-C. Folding analysis was carried out by RNAfold web server (http://rna.tbi.univie.ac.at/) (Gruber A.R. et al. 2008, Nucleic Acids Res., doi:10.1093/nar/gkn188) using DNA parameter settings at 37oC and 0.2 M salt. **B**, Mountain plot analysis of ssDNA folding for RRS-G4, RRS-C, and two strands of a 222 bp control DNA fragment from pEAW1307 plasmid. The observed high similarity between the MFE secondary structure (red line), the Centroid structure (blue line), and pair probabilities (Pf- green line) for RRS-G4 and RRS-C strongly indicate a reliable prediction of secondary structure for these sequences. Significant differences between MFE, Centroid and Pf lines for control sequences suggest that they are not likely to form stable secondary structures when present as ssDNA.

**Extended Data Fig. 2.**
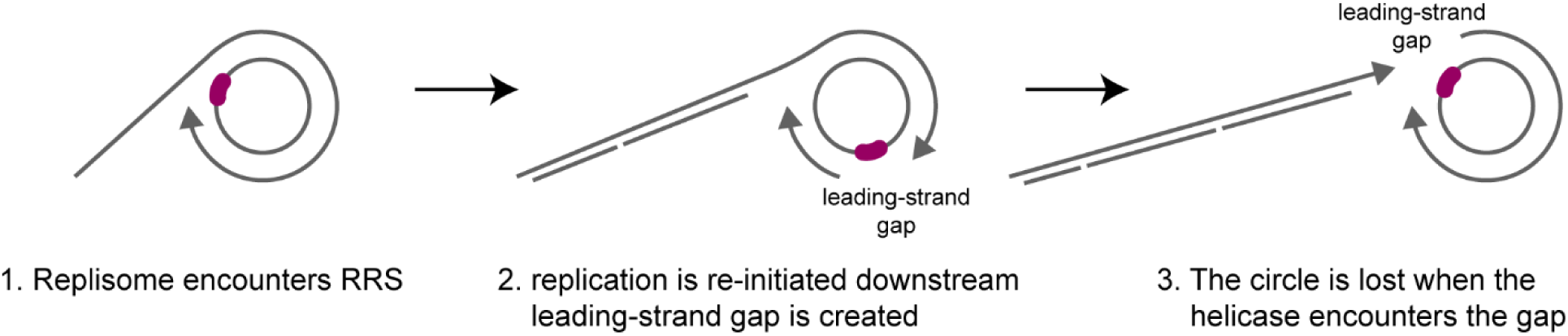
Formation of leading-strand gaps would terminate rolling-circle replication. Schematic representation of what would happen in the rolling-circle assay if a leading-strand gap were generated. If the leading-strand polymerase were to encounter a fully folded RRS-G4 (1), and leading-strand synthesis were reinitiated downstream, generating a leading-strand gap (2), the replisome would eventually lose the circular template upon helicase encounter with the gap, resulting in replication termination. We do not observe this.

**Extended Data Fig. 3.**
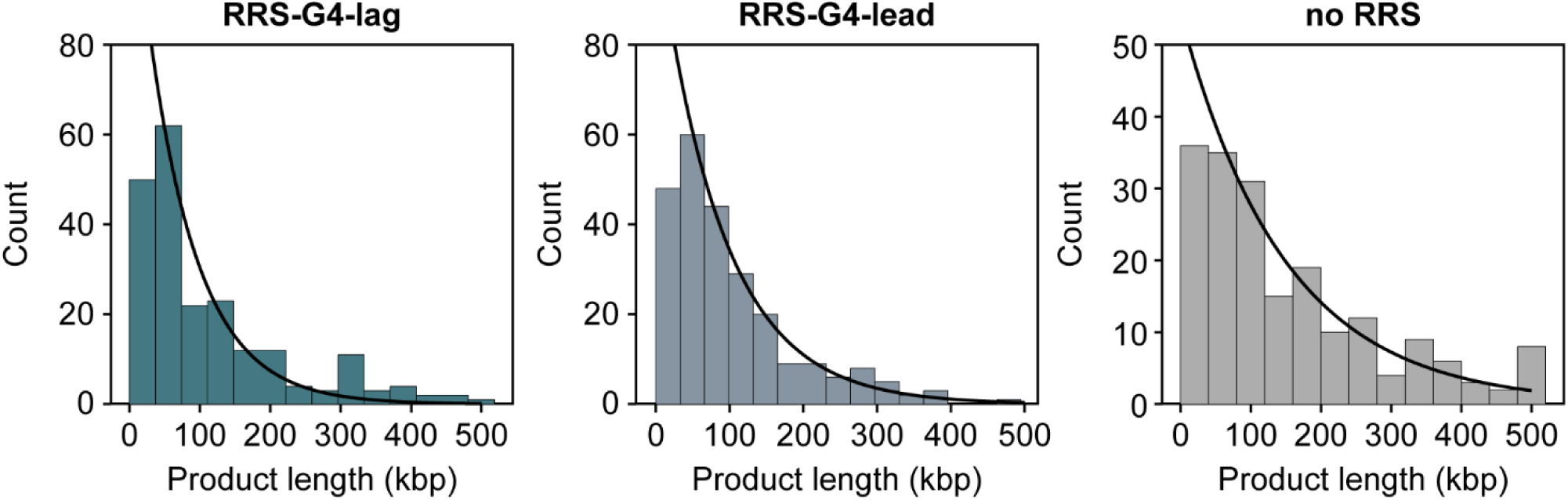
Length distributions. Distribution of the final product length of the replicated RRS-G4-lag (teal), RRS-G4-lead (blue) and control templates (grey). Each dataset is fit with a single-exponential decay function (black line), giving characteristic lengths of 71 ± 8 pix (n = 245) for RRS-G4-lag, 88 ± 9 pix (n = 211) for RRS-G4-lead, and 149 ± 12 pix (n = 191) for the control template.

**Extended Data Fig. 4.**
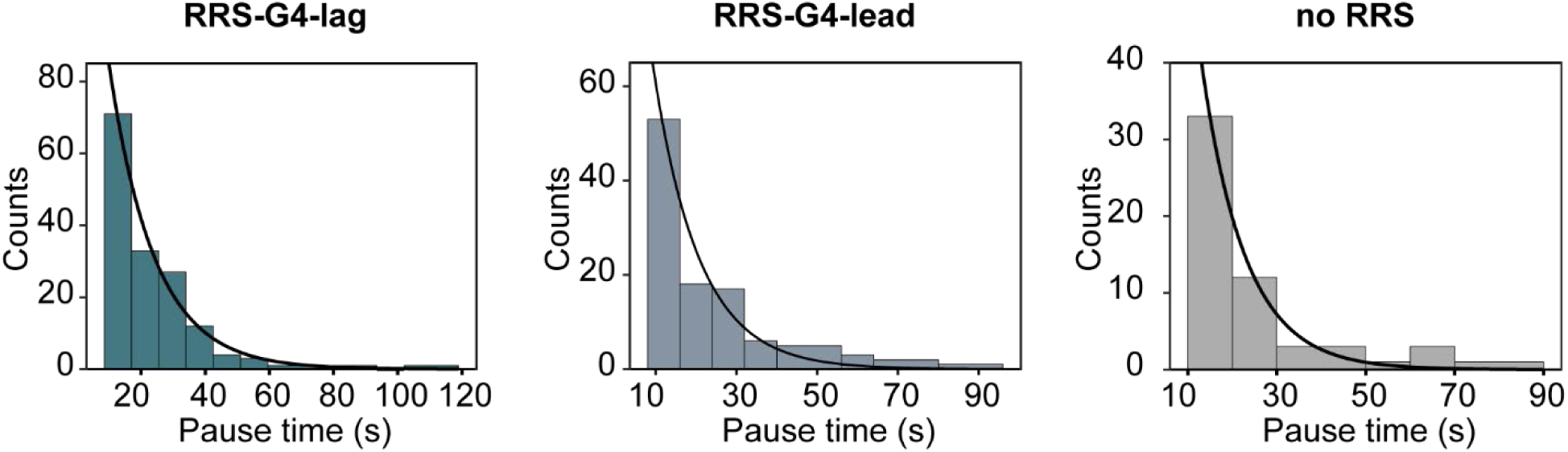
Pause duration distributions. Distribution of the pause times extracted from the rate histograms, with the replisome deemed to be paused when replicating less than 100 bp/s. The mean pause times are shown for the RRS-G4-lag (teal), RRS-G4-lead (blue) and control templates (grey). Each dataset was fit with a single-exponential decay (black line). The mean pause durations (± SEM) are 14.1 ± 1.1 s for RRS-G4-lag (n = 156), 11.4 ± 1.1 s for RRS-G4-lead (n = 113), and 9.87 ± 1.3 s for the control template (n = 57).

**Extended Data Fig. 5.**
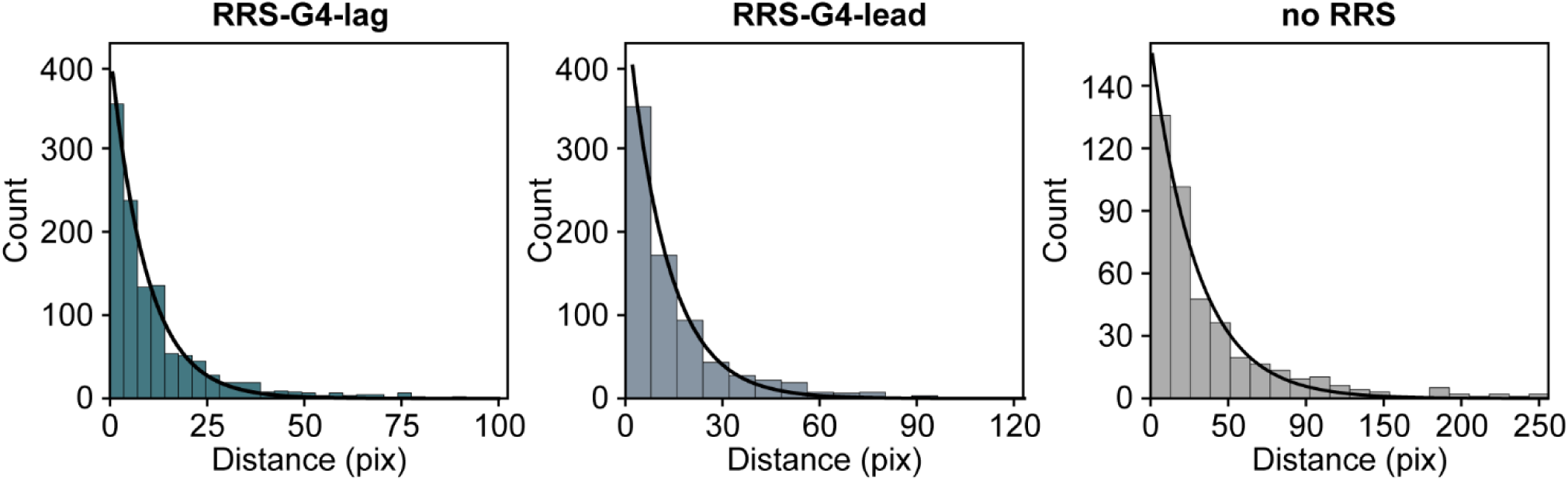
Distributions of the distance between the gaps. Distribution of the gap distances in the replicated RRS-G4-lag (teal), RRS-G4-lead (blue) and control templates (grey). Each dataset is fit with a single-exponential decay function (black line). The characteristic distances are 4.54 ± 0.14 kbp (n = 1,036) for RRS-G4-lag, 6.40 ± 0.24 kbp (n = 716) for RRS-G4-lead, and 15.62 ± 0.78 kbp (n = 403) for the control template.

**Extended Data Fig. 6.**
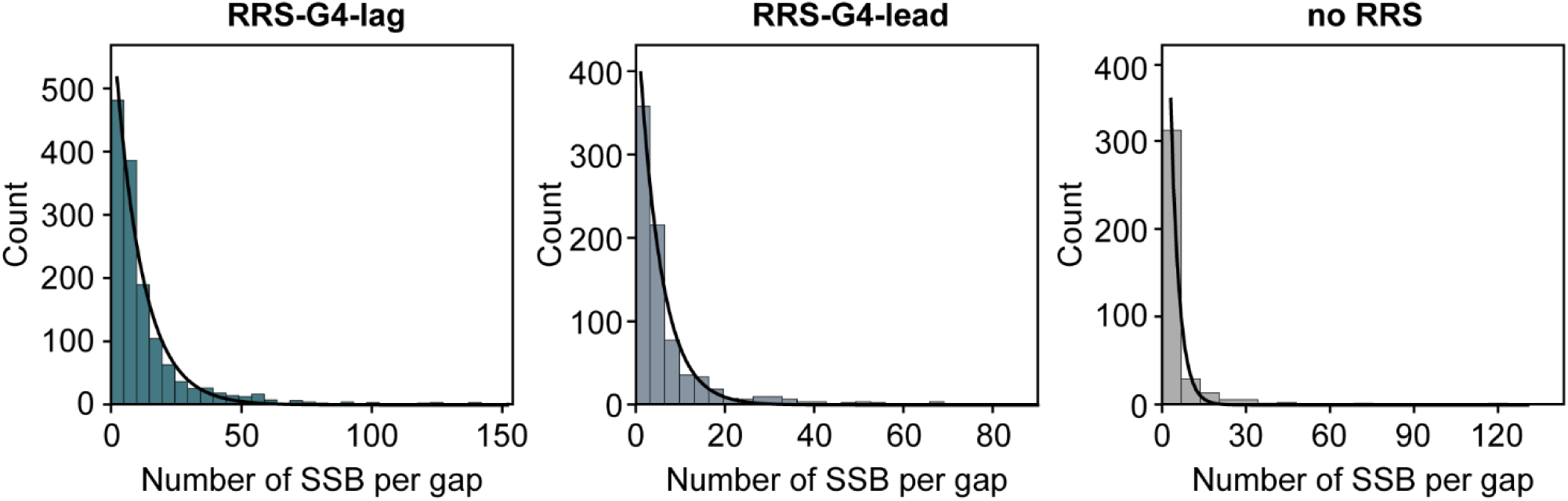
Distributions of the number of SSBs bound in the gaps. Distribution of the gap sizes in the replicated RRS-G4-lag (teal), RRS-G4-lead (blue) and control templates (grey). Each dataset is fit with a single-exponential decay (black line). The mean gap sizes (± SEM) are 698.4 ± 18.2 nt for RRS-G4-lag (n = 1,415), 328.3 ± 11.7 nt for RRS-G4-lead (n = 810), and 197.6 ± 10.4 nt (n = 386) for the control template.

**Extended Data Fig. 7.**
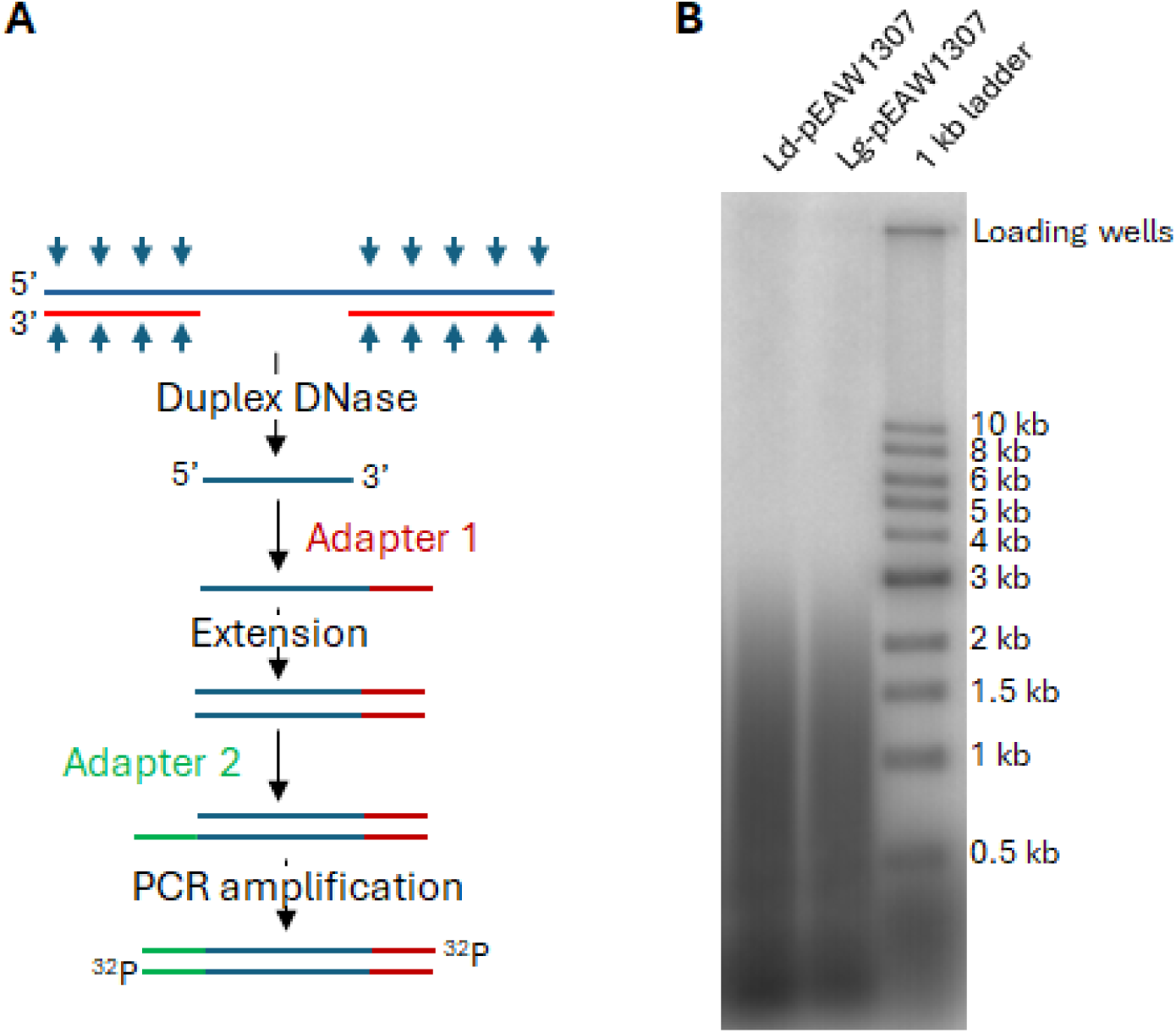
Size distribution of ssDNA gaps in replicated pEAW1307. **A**, Sketch of the experiment to digest replicated dsDNA and amplification of ssDNA gap fragments for size measurement. **B**, Distribution of ^32^P labelled DNA fragments amplified from ssDNA fragments after duplex DNAse treatment of replicated RRS-G4-lag and RRS-G4-lead. DNA were separated on a 1% agarose gel and visualised by phosphor-imaging.

**Extended Data Table 1.**
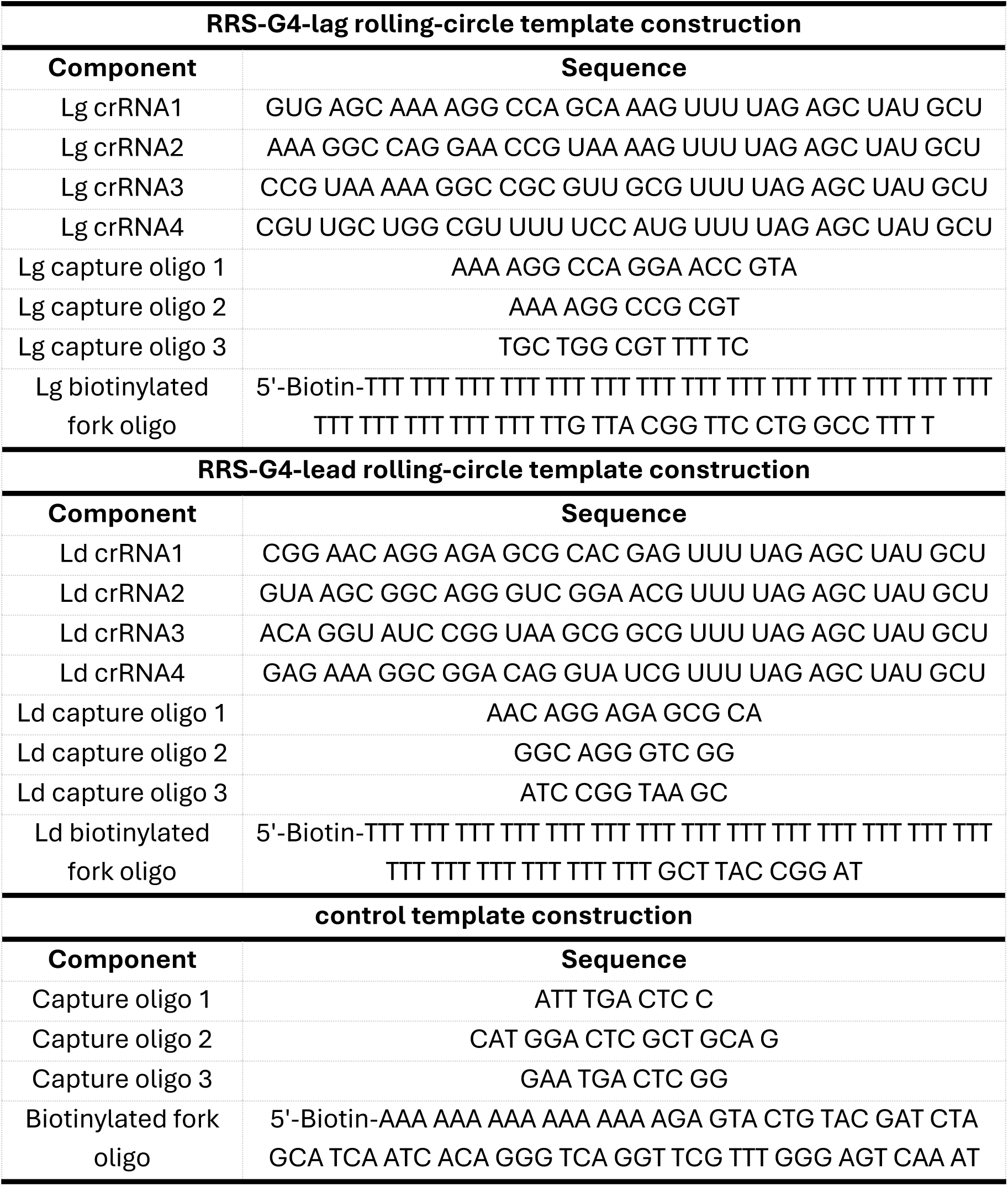
Oligos used in this study for template constructions.

